# Sex- and age-differences in cellular hallmarks of aging in a species with female-biased longevity and environmental sex determination

**DOI:** 10.64898/2025.12.10.693506

**Authors:** Jamie R. Marks, Fredric J. Janzen, Beth A. Reinke, Elizabeth A. Addis, Olumide Adesioye, Samantha Bock, Morgan Clark, Claudia Crowther, Luke A. Hoekstra, Jessica Judson, Caleb Krueger, Alexandria P. Sills, IISAGE Consortium, Anne M. Bronikowski

## Abstract

Cellular hallmarks of aging have been discovered and characterized in a number of model species for studying aging biology - such as humans, mice, fruit flies, and nematodes. Whether these canonical age-related changes to cellular physiology are present across diverse species that have variable rates of demographic aging remains less studied. Here, we tested whether several ubiquitous cellular hallmarks of aging – mitochondrial function, reactive oxygen species generation, and inducible DNA damage – change with age and in a sex-dependent manner in a species with indeterminate growth and reproduction (painted turtles, *Chrysemys picta*). A further feature of their biology that recommends them for an ecological model of vertebrate aging is their female-biased longevity, despite an absence of genotypic sex determination. Thus lifespan and aging may be reliable features of sex-specific life-histories. We measured aspects of mitochondrial health (cellular basal, maximal, and spare oxygen consumption rates), cellular levels of reactive oxygen species, and aspects of DNA damage and repair from exposure to UVB. We used these measures across several physiological axes as proxies for age-related physiological dysfunction. We further assessed our measures across several populations of painted turtles. We found that sex explained the largest proportion of variation, with males differing from females in mitochondrial function, reactive oxygen species production, and inducible DNA damage. In several cases, age significantly interacted with sex, but the effect size was small relative to sex alone. Thus, we found that sex, rather than age or size, was a consistent predictor of cellular aging physiological in this species with where females live longer and age slower.

## Introduction

Organismal aging – the age-related decline in cellular efficiency and organismal resiliency – is common across the animal tree of life and often manifests at the population level as accelerating age-specific mortality trajectories across adult lifespans (Reinke et al. 2022; Murthy and Ram 2015; Flatt and Partridge 2018). As a field, we have an established understanding of the evolutionary forces that can give rise to aging, which center on age-specific mutation-selection balances (Jové et al. 2023; reviewed in Promislow and Bronikowski, 2006) and have identified generalizable cellular hallmarks of aging (López-Otín 2013, 2023). However, whether and how cellular hallmarks of aging underly organismal aging broadly across taxa remains less understood. Specifically, there are 12 central cellular hallmarks of aging that span genome and cellular phenotypes (e.g., genomic instability, mitochondrial dysfunction, inflammation, etc.; López-Otín et al., 2023). Assessing these hallmarks in non-traditional species provides a unique opportunity to test their generalizability across species with variation in life histories, thermoregulatory modes, and sex-determining mechanisms (Koch et al. 2021, Biga et al. 2025). Accordingly, in this study we address the paucity of knowledge of cellular mechanisms of aging in diverse, long-lived species (Bronikowski et al. 2011; Harman et al. 2021; Holtze et al. 2021) in an effort to form a more integrative understanding of mechanisms of aging, how they evolve, and uncover their ecological drivers.

In this study, we focus on cellular hallmarks of aging that have a rich history of study in evolutionary physiology: mitochondrial physiology and DNA repair. With advancing age, decreasing mitochondrial efficiency (ATP produced per oxygen consumed), decreasing mitochondrial spare respiratory potential, and increasing loss of electrons in the electron transport chain, which can form damaging reactive oxygen species, are all features of the oxidative stress/mitochondrial health mechanism of aging (Harman 1956, Robert et al. 2007; Guo et al. 2013; Chistiakov et al. 2013; Hood et al. 2018; Gangloff et al. 2020; Li et al. 2023). Whether declining mitochondrial function occurs with advancing age across species, and whether declines occur at the same rate between males and females and/or across populations within species remain understudied (but see Robert et al. 2007; Rogell et al., 2014; Stier et al. 2017; Koch and Hill, 2018; Gangloff et al. 2020; Stier et al. 2022). Simultaneously, there is a growing appreciation that changes in mitochondrial function over the adult lifespan likely reflect a response to direct cellular damage occurring at the genomic level (Son and Lee, 2021). Declining efficiency of DNA damage repair pathways is another key hallmark of aging that has been assessed in the two sexes (e.g., *Drosophila melanogaster*: Aw et al. 2017; Nagarajan-Radha et al. 2019; *Rattus norvegicus:* Guevara et al. 2011; *Homo sapiens:* Viña et al. 2003). This has been studied in mammals, such as humans and mice, where rates of DNA damage accumulation (i.e., somatic mutations) is higher in males in species with female-biased longevity (Broestl and Rubin, 2021; Cardano et al., 2022), but negligibly in long-lived species (but see Robert et al. 2007, Firsanov et al. 2025).

An important predictor of both expected lifespan and aging rates is biological sex (Bronikowski et al. 2022). Current theories about the ultimate causes of sex specific ageing implicate mutation accumulation differences due to sex chromosomes as a potential factor. When chromosomal differences are present at conception, hypotheses have been proposed that include the “toxic Y” and “unguarded X”, both of which invoke mutation accumulation dynamics specific to the presence of heteromorphic sex chromosomes (Dunford et al., 2017; Nguyen and Bachtrog, 2021). For example, certain genes on the X-chromosome implicated in tumor-suppression can ‘escape’ inactivation in females, providing protection against single mutations at these loci (Dunford et al., 2017). Additionally, for all sex-determining mechanisms – ranging across genotypic to environmental forms – a general hypothesis has been established that males should age faster if mitochondrial function is a general correlate of aging. The “mother’s curse” hypothesis is based on mitochondrial inheritance through maternal lines, with males devoid of selection against mitochondrial mutation accumulation across generations, thus negatively impacting male aging phenotypes within generations (Dowling and Adrian, 2019). Other mechanisms of sex-specific demographic aging and age-related deterioration, such as differences in genome architecture, have been examined in model species such as mice *(Mus musculus*; Yuan et al. 2011; Bresilla et al., 2022), fruit flies (*Drosophila melanogaster*; Hoffman et al., 2014; Brown et al., 2020), and/or nematodes (*Caenorhabditis elegans*; Olsen et al., 2006; Hotzi et al. 2018; reviewed in Bronikowski et al., 2022). These studies have provided insights into causes of sex differences in aging and are largely supportive of the ubiquitous nature of molecular mechanisms underlying aging (López-Otín et al. 2023) – or at least in species that have lent themselves to these types of experiments in the past. Gaining a deeper understanding of the mechanisms contributing to sex differences in aging and lifespan enhances the overall understanding of sex-specific life-histories.

The effect of unequal genome architecture between the sexes can disproportionately influence our understanding of sex-specific aging. Therefore, species that lack sex chromosomes provide a unique opportunity to study sex-specific differences without these confounding factors. Painted turtles (*Chrysemys picta*) are one such species – they lack sex chromosomes and instead having temperature-dependent sex determination (i.e., sex is determined by incubation temperature during the middle third stage of egg incubation, where higher nest temperatures lead to female biased clutches; Bull and Vogt, 1979; Janzen, 1994). These turtles also display sex-specific aging, with males aging more rapidly than females (Congdon et al., 2013; Bronikowski et al., 2022). Herein, we measure mitochondrial function by assessing basal and maximal cellular oxygen consumption rates as well as spare respiratory potential in white blood cells from whole blood collections – a non-terminal sampling method. We further measure intracellular levels of two reactive oxygen species (mitochondrial superoxide and intracellular hydrogen peroxide). As measures of a second hallmark of aging, we assess baseline and inducible DNA damage, and repair efficiency in erythrocytes from whole blood –for a homogeneous cell population –from the same individuals.

We test the hypothesis that mitochondrial health is affected by age, and we make the specific predictions that mitochondrial health declines with age, and inducible DNA damage increases with advancing age in painted turtles. Whether these hallmarks decline with age in turtles is of particular interest given that the Testudines clade is characterized by exceptionally slow demographic aging and concomitantly long lifespans when corrected for body size (Reinke et al., 2022). We further test the hypothesis that age-specific changes in these hallmarks follow sex-specific patterns and predict that these changes are more pronounced in males, the faster-aging sex. This is also of particular importance, given that sex is determined through thermally-sensitive gene expression during the critical developmental stage (Rhen and Schroeder, 2017). In accordance with the oxidative stress hypothesis, we predict that measures of mitochondrial health would be greater in young adults for both sexes and deteriorate more quickly in males. We also predict that inducible DNA damage would be greater in older animals, particularly males, while repair capabilities would be greater within younger animals, particularly young females. Finally, we test whether the influences of sex and age on mitochondrial function and DNA repair varies across three geographically-distant populations of painted turtles, to assess whether age-by-sex interactions are fixed versus plastic across space (see Judson et al., 2020 which tests genetic diversity across the three painted turtle populations), which lends insight into the evolution of these phenotypes.

## Methods

### Study species

Painted turtles *(Chrysemys picta)* have longer than expected lifespans based on their body size, coupled with slow rates of accelerating mortality across adulthood (Congdon et al. 2013; Warner et al., 2016; Reinke et al., 2022). Gonadal sex is determined permanently by thermally-sensitive cascades of gene expression during the middle third of incubation. In this species, females are produced at warmer temperatures with males at cooler temperatures. The pivotal temperature for expected 1:1 sex ratio is 28°C on average across its wide geographic range (Carter et al. 2019) and, accordingly, maternal behavior ensures relatively similar nest thermal environments among populations despite divergent local climates (Carter et al., 2019; Bodensteiner et al. 2023). Despite the lack of genotypic sex determination, painted turtles have sex-specific life histories, in which males mature earlier and age faster than females. Males mature between 3-5 years of age and, in our main study site in Illinois, can live to 25 years. Females mature between 5-7 years and can live 40+ years (Schwanz et al., 2010; Spencer and Janzen, 2010; Hoekstra et al., 2018; Bronikowski et al. 2022). The differences in lifespan are underlain by differences in mortality acceleration across adulthood. In Illinois, female mortality rate doubling time is 35 years, whereas that for adult males is 10 years (Bronikowski et al. 2022). Painted turtles also exhibit indeterminate growth and fecundity, though growth slows in later adulthood (Congdon et al. 2013; Hoekstra et al. 2018). Because of this, shell length (plastron length for painted turtles) is tightly correlated with age in general, particularly during the early portion of the lifespan before growth slows appreciably after maturity.

### Study populations

We included individuals from three populations of painted turtles across their range from Illinois, Wisconsin, and Idaho. The Illinois population resides in Carroll County (42.0386° N, −89.9627° W) on the backwaters of the Mississippi River and has been the site of mark-recapture study for 36 years (Warner et al., 2016; Delaney et al., 2021). Due to this sustained study, a large proportion of animals captured each year are of known age based on initial captures at small size and discernable growth annuli. New animals are assigned an estimated age based either on plastron scute annuli or on published von Bertalanffy growth models for this population (Hoekstra et al., 2018; see Judson et al., 2020). Because age is known in the Illinois painted turtles, they are the focal population for age-based analyses. The Wisconsin population resides in Lac Courte Oreilles in Sawyer County (45.8889° N, −91.4405° W) and has been the subject of a mark-recapture study for 15 years (Reinke et al., 2020). The Idaho turtle population is found at Round Lake in Bonner County (48.1610° N, −116.6369° W) and has been studied intermittently for the past 12 years (Delaney et al. 2017; Carter et al. 2019) and is near the northern range limit of painted turtles (Holman and Andrews, 1994). Because age is not as well-characterized in the Wisconsin and Idaho populations, our analyses across the three populations used plastron length as a proxy for age (see *Sample Collection* below). Despite distinct population boundaries and disparate sampling locations, painted turtles across their geographic range exhibit low amounts of genetic differentiation (e.g., Idaho-Illinois Fst = 0.16; range across the species distribution = 0.04 – 0.20, excluding a relict population in New Mexico) and remarkable genomic homogeneity relative to other species at similar distances (Judson et al. 2024).

### Sample collection

In May through June of 2023 and 2024, females and males were trapped using fyke nets, baited aquatic traps, dip nets, or hand-sampled while on land. For each turtle, we noted recapture status, and measured morphometrics (weight, lengths and widths of carapace and plastron), as well the sex of each animal and any unique morphological features. From each animal, we collected 1 mL of whole blood in a heparinized insulin syringe (28 gauge) from the caudal vein. One drop of blood from the syringe was used to make a blood smear. These slides were dipped in methanol for one minute and transported to the lab for staining and determining white blood cell differentials (blood collection and smears described in detail in Holden et al. 2022). Remaining blood was mixed 1:1 with sterile medium (a DMEM-based medium supplemented with heparin and HEPES). Samples were kept cold and transported overnight on ice to the W. K. Kellogg Biological Station. Upon arrival, we immediately isolated peripheral blool mononuclear leukocytes (primarily lymphocytes and monocytes), and red blood cells (erythrocytes). The limited PBMC were used for cellular metabolism studies, whereas the more abundant homogeneous erythrocytes were used for the assays of reactive oxygene species and DNA repair. For this analysis, our sample sizes were N=67 from Illinois (24 females, 43 males); N=32 from Wisconsin (16 females, 16 males); and N=47 from Idaho (30 females, 17 males).

### Cellular oxygen consumption rate

Following guidelines established by Brand and Nicholls (2011), upon arrival to the laboratory, our workflow included isolation of white blood cells comprised primarily of lymphocytes and monocytes (analogous to isolating peripheral blood mononuclear cells – PBMCs – in species lacking nucleated red blood cells) with density gradient cell separation medium (Sigma Aldrich Histopaque; Cat. No. 10771); plating the cells in Agilent XFe24 cell culture plates; and running the Agilent mitochondrial stress test protocol (Cat. No. 103015-100) optimized for painted turtle leukocytes on an Agilent extracellular flux analyzer (XFe24). Leukocytes were plated at 1 million cells per well in Agilent DMEM medium (Cat. No. 103575-100) with at least two replicate wells per sample. When cell yields were limited, we seeded cells at 500,000 or greater and normalized to 1 million cells. In a separate pilot study, we ascertained that normalization to 1 million was valid for seeding densities greater than 450,000 cells per well. All runs and incubations were conducted at 28° C, which is a typical resting (non-basking) body temperature of painted turtles.

We used the manufacturer’s protocol to measure cellular oxygen consumption rates (OCR) in pmol/min during four phases of cellular respiration: (i) baseline OCR (in the presence of pyruvate as substrate, but no inhibitors); (ii) OCR during inhibition of ATP synthase; (iii) OCR during uncoupling of oxygen consumption from ATP synthesis; and (iv) OCR after further inhibiting both Complex 1 and Complex 3 of the electron transport chain. For each of these four phases, we took three measurements, and checked for acceptable variance within and among replicates (coefficient of variation less than 0.20). The output files for the assays were visually inspected for any failed wells (i.e., failed injections, occasional bubbles in the well). Then, we computed basal OCR [(i)-(iv)] and maximum OCR [(iii)-(iv)]. We computed spare respiratory potential as the difference between the maximum OCR and basal OCR. Our dependent variables were Basal OCR, maximum “Max” OCR, and spare respiratory potential (“Spare” hereafter). All raw data were in pmol/min per 1 million cells. In a few cases where we did not have this quantify of cells for triplicates, and results were normalized to 1 million cells.

### Leukocyte Differentials

To understand whether sexes and ages differed in leukocyte differentials, and thus potentially causing variation due to differences in cell populations, we conducted white blood cell differenetials from blood smears. Blood smears were made by depositing a drop (~ 3 ul) of fresh blood onto a 75 mm x 38 mm slide. The drop of blood was then spread across the slide with the edge of a secondary slide. Slides were dried, fixed with methanol, and later stained with Wright-Giemsa. Slides were stored in slide boxes until they were scored. Four types of leukocytes were counted: heterophils, lymphocytes, basophils, and monocytes. One hundred leukocytes per slide were counted and the cell types were recorded (Palacios et al., 2020; Holden et al., 2022). We analyzed whether populations and sexes differed in their proportions of different types of leukocytes because PBMCs are a heterogeneous population. We were particularly interested in the effect of size or age because white blood cell composition can change with age and could lead to incorrect interpretations of age-related results if, in fact, cell composition is changed with age. PBMCs are primarily lymphocytes and monocytes, but may also have additional cell types (such as basophils and heterophils) that can influence PBMC yields, and can be indicative of stress and inflammation.

### Reactive Oxygen Species

For the Illinois (known-age) painted turtles, we measured levels of mitochondrial superoxide (O_2_^−^) and intracellular hydrogen peroxide (H_2_O_2_) in red blood cells immediately post-isolation using MitoSOX® red (for O_2_^−^) (Invitrogen M36008) and CellROX® green (for (H_2_O_2_) (Invitrogen C10444). Both dyes exhibit fluorescence upon oxidation (Escada-Rebelo et al., 2020). Briefly, after PBMCs were isolated, we collected 50 µL of packed RBCs and resuspended them into 2 mL of DMEM medium and performed a cell count with trypan blue. We placed an appropriate volume of this cell suspension for 1.8 million cells in 300 µL DMEM with a final concentration of 5 µM MitoSOX® and CellROX®. Cells were incubated at 28°C for 30 mins. Cells were then washed twice with centrifugation and were resuspended in 300 µL of PBS. Three replicates of 600,000 cells/well, were plated in 100 µL in a Black 96 well optical plate. Three wells of PBS served as a negative control on each plate.

Plates were read using the BioTek Cytation 5 Cell Imaging Multimode (Agilent) plate reader and the Gen5 Image+ software. Incubation temperature during plate reading was 28°C, and a slow double orbital shake step was added for 5 minutes. Fluorescence was measured at Excitation/Emission wavelengths of 485/520 for CellROX® and 500/585 nm for MitoSOX® and Relative Fluorescence Units (RFUs) recorded. For a few individuals with limited cell yield, we plated fewer than 600,000 cells per well and normalized RFU to 600,000 cells.

### Single Cell “Comet” Alkaline Assay of DNA damage

We conducted the alkaline single-cell gel electrophoresis assay (Singh et al., 1988; aka “comet” assay), which we have used previously to obtain estimates of baseline DNA damage, inducible DNA damage upon treating cells with a genotoxic treatment, and DNA repair efficiency by allowing damaged cells time to repair (Robert and Bronikowski, 2010; Schwanz et al., 2011; Schwartz and Bronikowski, 2013). Red blood cells (erythrocytes) were obtained from the bottom layer of cells after centrifugation on the Histopaque density gradient (see above). Cells were resuspended in DMEM at a concentration of 1×10^5/mL. 5 uL of this mixture (i.e., 500 cells) were mixed with 50 uL low-melt agarose for each treatment (baseline, UV-induced damage, repair). This mixture of 500 cells in 50 uL LMA was spread evenly on a Comet slide according to manufacturer protocols (BioTechne, St Paul, MN). Baseline damage (“B” treatment) cells were lysed immediately; UV-damage (“D” treatment) cells were subjected to 5 mins of exposure to UV (312nm) on a transilluminator; Repair-damage (“R” treatment) cells were UV-damaged as above and allowed 10 mins at 28°C to repair DNA damage. For both Idaho and Wisconsin, freshly isolated cells were used to conduct the comet assay on the day of arrival of whole blood. For Illinois, because we used red blood cells for ROS measures, we followed our standard slow-freeze protocol in DMEM+DMSO and removed these live cells from a liquid nitrogen vapor layer one month post-freeze to conduct the comet assay. “Comet” slides from Idaho and Wisconsin were imaged using OpenComet ImageJ add-in (Gyori et al., 2021) and slides from Illinois were imaged with Comet Assay IV (version 4.3.2, Instem, Pennsylvania US). For all individuals, up to 50 comets were quantified per treatment. Both tools measure the proportion of DNA in the tail region of the comet versus the head region of the comet. Multiple slides were read in both software programs for cross-validation. From the resulting data for each turtle, we removed statistical outliers outside of 1.5 multiplied by the interquartile range and ensured our coefficients of variation were less than 0.20. We calculated inducible damage, defined as the proportional change between damaged cell comet tails and baseline cell comet tails ((D-B)/D), and repair efficiency, defined as the change between damaged cells and cells allowed to repair, relative to baseline damage ((D-R)/B).

### Statistics

We undertook two analysis workflows – the first for Illinois animals who were of known age based on recapture data, the second for animals from all three populations, including two populations where age was not known and, therefore, plastron length was used as a proxy for age and size-based analyses were performed.

#### Known-age analyses (Illinois)

Our overarching goal was to test for the effects of sex, age, and their interaction on our dependent variables. Because age differs between males and females, we Z-transformed observed age to a mean of zero and unit variation within each sex, which removed its association with sex, and allowed for an independent comparison of relatively old females to relatively old males, and so on [(Mean Yrs, Range), Females (n = 22): 15.2, 7 – 36; Males (n = 38): 7.5, 3 – 22; F_1,58_ = 15.85, *Pr* = 0.0002)]. We used this variable “zAge” in our age-specific analyses for Illinois painted turtles.

#### Size-based analyses (all populations)

Preliminary analyses indicated that females had significantly greater plastron length (PL) than males in each population (see Table S1). Thus, we again performed a Z-transformation of PL for each population-and-sex grouping (i.e., six groups of animals: three populations x two sexes). This approach allowed us to compare relatively larger/older animals to relatively smaller/younger animals across populations and sexes. We used this variable “zPL” in our size-specific analyses across all three populations.

#### Dependent variables and transformations

We modeled variation in cellular metabolism as a proxy for mitochondrial function (Basal OCR, Max OCR, Spare); levels of mitochondrial superoxide and intracellular hydrogen peroxide as proxies for mitochondrial health; and UV-inducible damage and repair efficiency as proxies for the DNA-damage-repair response. These seven variables together are the focal variables of interest. In addition, we analyzed leukocyte counts to ascertain whether any changes we identified in cellular metabolism were attributable to differences in cell populations (and not age, sex, or size).

We ran Shapiro-Wilk’s tests to test for normality. To improve normality, we log_10_ transformed Basal OCR, Max OCR, and Spare, as is customary with oxygen consumption data. We natural-log (ln or log_e_) transformed normalized mitochondrial superoxide (O_2_^−^), intracellular hydrogen peroxide (H_2_O_2_) RFU, and leukocyte counts. Because our measures of inducible DNA damage and repair efficiency were proportions that could range above 1.0, we logit transformed these data, along with their underlying proportions (i.e., proportion DNA in comet tails under baseline (B), UV-damaged (D), and damage + repair (R)). Once all variables met normality assumptions, we removed from one to three outliers for each assay based on values outside 1.5 times the interquartile range.

For our age-based analyses, conducted only for the Illinois population of painted turtles, we conducted ANCOVA and used Type III sums of squares due to imbalance of sample sizes, using the following model:

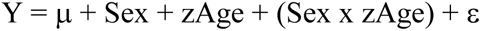

where Sex is male or female, and zAge is sex-specific z-transformed age. This model was used to analyze log_10_(Basal OCR), log_10_(Max OCR), log_10_(Spare), ln(O_2_^−^ induced fluorescence), ln(H_2_O_2_ induced fluorescence), and logit(inducible damage). Due to the small sample size, and intentional selection of the oldest and youngest males and females for the Illinois known-age DNA-damage assay, we used Categorical Age (i.e., Aged versus Young adults) for the analysis of inducible damage.

For our multi-population size-based analyses, we conducted an analysis of covariance using the following model including Population (“Pop”: IL, WI, ID), Sex (male, female), and the continuous covariate zPL.

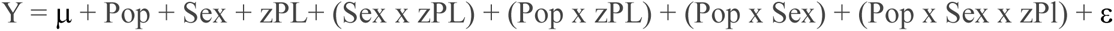

where the third order interaction is the main term of interest to test for population differences in sex-by-size relationships. Models were run in SAS 9.4 (Cary, North Carolina) with Proc GLM and type III sums of squares.

## Results

### Age-based cellular aging hallmarks (Illinois only)

#### Cellular Metabolism (mitochondrial function)

Males had substantially higher cellular Basal OCR, Max OCR, and Spare than females (Table 1, Fig. 1). Males had marginally increasing Basal OCR with age relative to females, and females had marginally increasing Spare with age relative to males. Notwithstanding, this interaction explained only 4% of variation, with sex explaining more than 20% for each dependent variable.

**Table 1.** Analysis of variance of log_10_ cellular oxygen consumption rates (pmol/min): Basal OCR, Max OCR, and Spare in the Illinois population of painted turtles. Effect is the proportion of variation explained for explanatory variables with Pr < 0.15. NS is Pr > 0.20. zAge is z-transformed age in years to remove its association with Sex. Sex is male and female. See text for details of models.

|  | **Basal OCR (log<sub>10</sub>)** |  | **Max OCR (log<sub>10</sub>)** |  | **Spare (log<sub>10</sub>)** |  |
| --- | --- | --- | --- | --- | --- | --- |
| **Source of Variation** | **F<sub>df1, df2</sub>**<br>**(*Pr* > F)** | **Effect** | **F<sub>df1, df2</sub>**<br>**(*Pr* > F)** | **Effect** | **F<sub>df1, df2</sub>**<br>**(*Pr* > F)** | **Effect** |
| Sex | 13.59<sub>1,54</sub><br>(0.0006) | 0.20<br>M > F<br>Fig. 1 | 24.15<sub>1,48</sub><br>($<0.0001$) | 0.31<br>M > F | 15.67<sub>1,47</sub><br>(0.0003) | 0.24<br>M > F |
| zAge | 0.10<sub>1,54</sub><br>(NS) | -- | 2.26<sub>1,48</sub><br>(0.14) | 0.03<br>Aged ><br>Young | 2.69<sub>1,47</sub><br>(0.11) | 0.04<br>Aged ><br>Young |
| Sex x zAge | 2.81<sub>1,54</sub><br>(0.09) | 0.04<br>F: Aged <<br>Young<br>M: -- | 1.36<sub>1,48</sub><br>(NS) | -- | 2.33<sub>1,42</sub><br>(0.13) | 0.04<br>F: Aged ><br>Young<br>M: -- |

**Figure 1.**
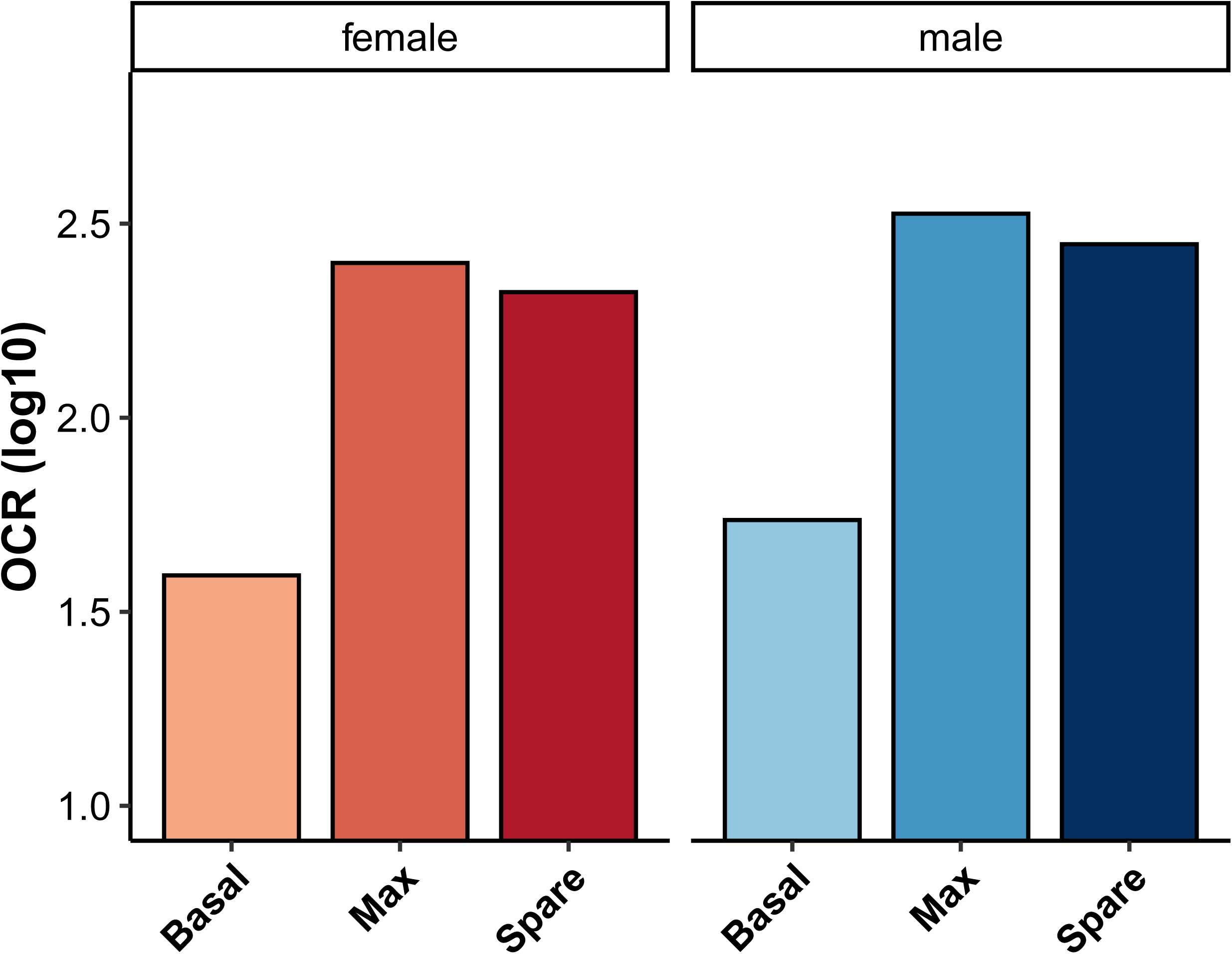
Effect of sex on cellular OCR (oxygen consumption rate in pmol/min/1 million cells) for basal, maximal, and spare respiratory potential for Illinois painted turtles. Data are LSmeans from a model accounting for zAge (see Table 1 for statistical output) with log_10_ transformed dependent variables. Male values were greater than female values for all three measures of cellular OCR.

#### Cellular levels of reactive oxygen species

Sex and zAge significantly interacted to affect both mitochondrial superoxide (O_2_-) and intracellular hydrogen peroxide (H_2_O_2_). Males had lower levels of both O_2_- and H_2_O_2_ at older ages, whereas female levels did not change with age (Table 2, Fig. 2). Additionally, like cellular metabolism, male cellular ROS was higher than female cellular ROS, regardless of age.

**Figure 2.**
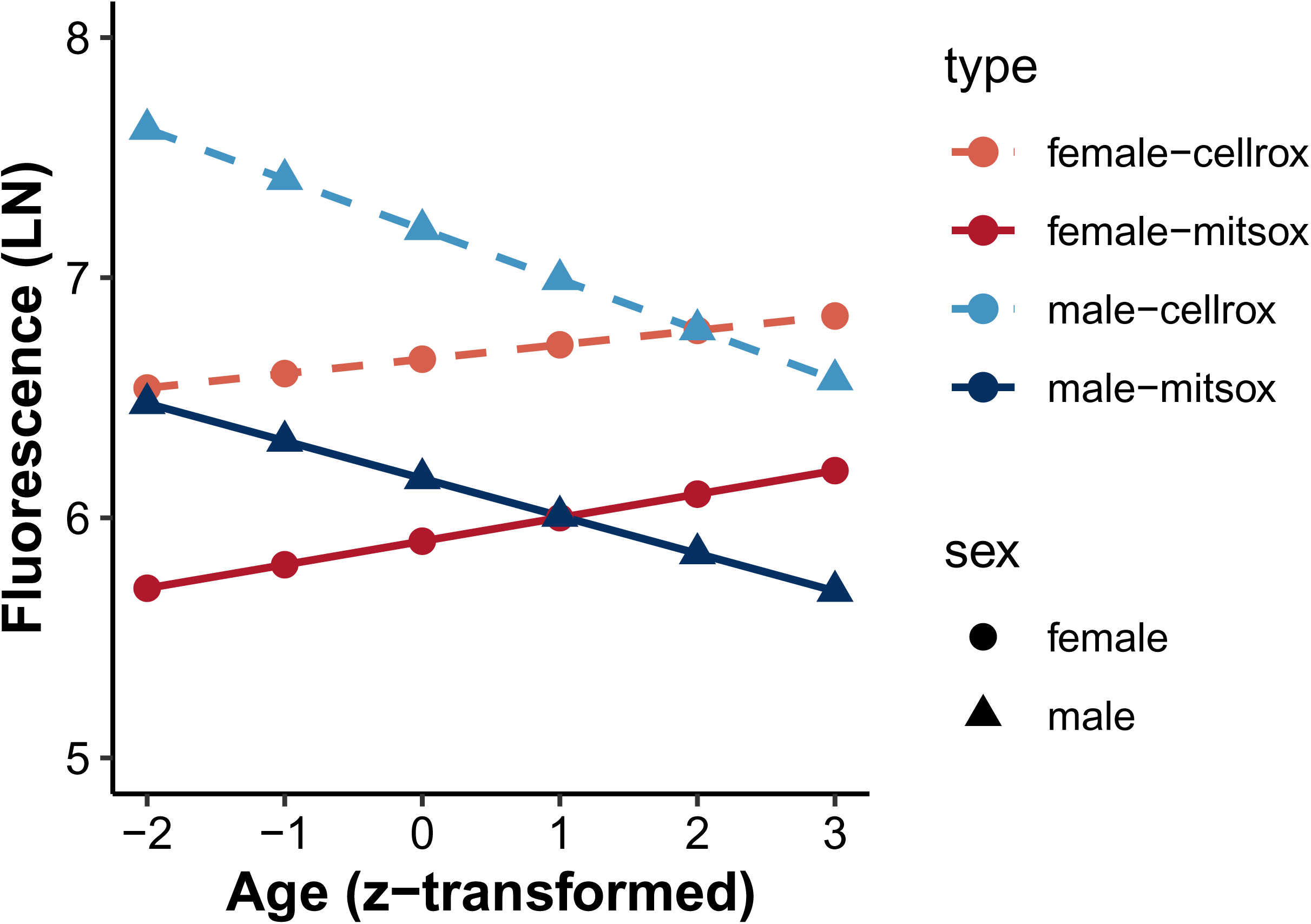
Interaction effect of (Sex x zAge) on mitochondrial superoxide RFU (solid lines) and total cellular hydrogen peroxide RFU (dashed lines) for Illinois painted turtles (see Table 2 for statistical output). Illinois Females (circles, solid line) have no change in mitochondrial superoxide with increasing age (slope = 0.10, *Pr* = NS); males (triangles, solid line) have decreasing mitochondrial superoxide with age (slope = −0.16, *Pr* = 0.03). For cellular hydrogen peroxide, females (circles dashed line) have no change in cellular H_2_O_2_ with increasing age (slope = 0.06, *Pr* = NS); males (triangles, dashed line) have decreasing cellular H_2_O_2_ with age (slope = −0.21, *Pr* = 0.01). Data are LSmeans estimated at six values of zAge from a model accounting for zAge (see Table 2 for statistical output) ln (i.e., log*_e_*) transformed dependent variables.

**Table 2.** Analysis of Variance of natural log of normalized RFU for mitochondrial superoxide (O_2_-) (measured by MitoSOX® red), total cellular H2O2 (measured by CellROX® green), and logit of % inducible DNA damage in the Illinois population of painted turtles. Effect is the proportion of variance explained for effects with Pr <0.15. NS is Pr > 0.20. Sex is male and female; zAge is z-transformed age in years; Categorical Age is Aged and Young adults.

|  | **Mitochondrial**<br>**Superoxide(O<sub>2</sub><sup>-</sup>) (ln)** |  | **Cellular ROS (H<sub>2</sub>O<sub>2</sub>)**<br>**(ln)** |  | **UV-induced damage**<br>**(logit)** |  |
| --- | --- | --- | --- | --- | --- | --- |
| **Source of**<br>**Variation** | **F<sub>df1, df2</sub>**<br>**(Pr > F)** | **Effect** | **F<sub>df1, df2</sub>**<br>**(Pr > F)** | **Effect** | **F<sub>df1, df2</sub>**<br>**(Pr > F)** | **Effect** |
| Sex | 4.31<sub>1,61</sub><br>0.04 | 0.06<br>M > F | 15.14<sub>1,61</sub><br>(0.0002) | 0.1<br>M > F | 5.21<sub>1,9</sub><br>(0.05) | 0.21<br>M > F |
| zAge | 0.22<sub>1,61</sub><br>(NS) | -- | 1.18<sub>1,61</sub><br>(NS) | -- | NA | NA |
| Sex x zAge | 4.14<sub>1,61</sub><br>(0.05) | 0.06<br>(Fig. 2a) | 3.78<sub>1,61</sub><br>(0.06) | 0.04<br>(Fig. 2a) | NA | NA |
| Age<br>Category | NA | NA | NA | NA | 2.47<sub>1,9</sub><br>(0.15) | -- |
| Sex x Age<br>Category | NA | NA | NA | NA | 5.80<sub>1,9</sub><br>(0.04) | 0.24<br>Young<br>Adult Males<br>> All Other<br>Groups |

#### Inducible DNA damage

We found that UVB-induced damage was higher in younger male turtles compared to younger females, and higher than in older animals of both sexes (Table 2). Erythrocytes of young male turtles (ca. 3-6 yrs) had greater levels of DNA damage (measured as %DNA in the comet tail versus the comet head) after exposure to UVB radiation than any other group (LS means of logit[(Damaged – Baseline)/Damaged] Young adult male (YM) > Aged adult male (AM) = Young adult female (YF) = Aged adult female (AY): 5.84 > 2.22 = 1.58 = 2.34. (The back-transformed values equate to roughly 98% inducible damage in young adult males versus a mean of 90% for all other age and sex categories).

### Cellular aging hallmarks across three populations

#### Cellular Metabolism (mitochondrial function)

Source population, sex, and size (z-transformed plastron length) significantly interacted to affect cellular basal oxygen consumption rate, maximal OCR and cellular spare respiratory potential. Importantly, however, there was no consistent pattern (such as all females increase basal oxygen consumption with advancing age, but at different rates; Table 3). Source population and sex interacted to significantly affect all basal, Maximal, and Spare in a uniform manner. In all cases, and in agreement with the age-based analyses above, males had higher Basal OCR, Max OCR, and Spare (Table 3; Fig. 3), but by different magnitudes.

**Figure 3.**
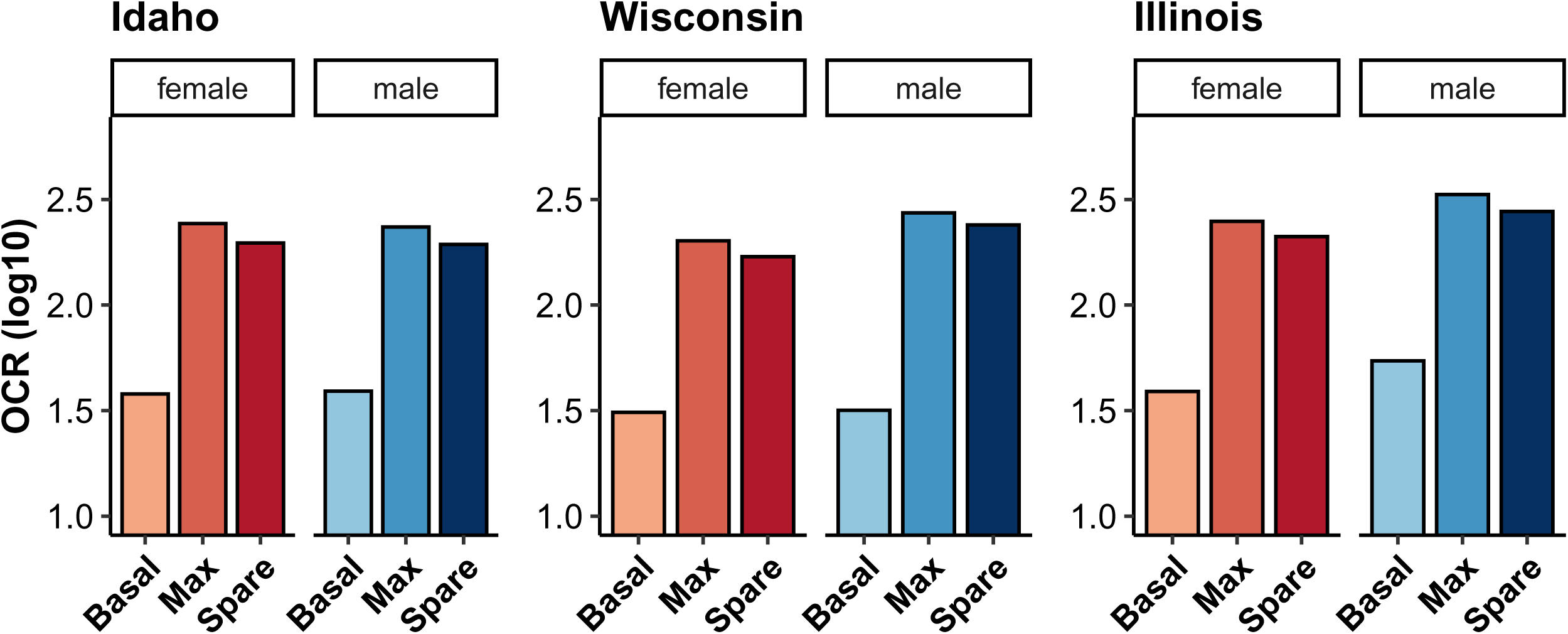
Least-square means for the interaction effect of Sex-by-Population for log_10_ cellular Basal OCR, Max OCR, and Spare. All variables differed between males (blue) and females (red) within populations, and by different magnitudes across populations. Males (blue) and females (red). See Table 3 the full models.

**Table 3.** Analysis of variance of log_10_ cellular oxygen consumption rates (pmol/min): Basal OCR, Max OCR, and Spare in three populations of painted turtles. Effect is the proportion of variation explained for explanatory variables with Pr < 0.15. NS is Pr > 0.20. zPl is z-transformed plastron length for each sex-by-population grouping to remove its association with both sex and population. Pop is population. See text for details of models.

|  | **Basal OCR (log<sub>10</sub>)** |  | **Max OCR (log<sub>10</sub>)** |  | **Spare (log<sub>10</sub>)** |  |
| --- | --- | --- | --- | --- | --- | --- |
| **Source of Variation** | **F<sub>df1, df2</sub>**<br>**(*Pr* > F)** | **Effect** | **F<sub>df1, df2</sub>**<br>**(*Pr* > F)** | **Effect** | **F<sub>df1, df2</sub>**<br>**(*Pr* > F)** | **Effect** |
| Pop | 14.76<sub>2,119</sub><br>($<0.0001$) | 0.17 | 6.26<sub>2,113</sub><br>(0.003) | 0.10 | 5.37<sub>2,110</sub><br>(0.006) | 0.07 |
| Sex | 5.31<sub>1,119</sub><br>(0.03) | 0.03<br>M > F | 11.38<sub>2,113</sub><br>(0.001) | 0.09<br>M > F | 10.58<sub>1,110</sub><br>(0.002) | 0.07<br>M > F |
| zPl | 1.87<sub>1,119</sub><br>(NS) | -- | 0.14<sub>1,113</sub><br>(NS) | -- | 0.70<sub>1,110</sub><br>(NS) | -- |
| Pop x Sex | 4.02<sub>2,119</sub><br>(0.02) | 0.05<br>(Fig. 3) | 4.08<sub>2,113</sub><br>(0.02) | 0.06<br>(Fig. 3) | 3.04<sub>2,110</sub><br>(0.05) | 0.04<br>(Fig. 3) |
| Pop x zPl | 0.39<sub>2,119</sub><br>(NS) | -- | 0.77<sub>2,113</sub><br>(NS) | -- | 0.74<sub>2,110</sub><br>(NS) | -- |
| Sex x zPl | 0.27<sub>1,119</sub><br>(NS) | -- | 3.37<sub>1,113</sub><br>(0.07) | -- | 2.09<sub>1,110</sub><br>(NS) | -- |
| Pop x Sex x zPl | 2.78<sub>2,119</sub><br>(0.06) | 0.03 | 2.34<sub>2,113</sub><br>(0.10) | 0.04 | 2.75<sub>2,110</sub><br>(0.07) | 0.04 |

#### Inducible DNA damage and repair efficiency

Both population and sex affected inducible DNA damage, with males having higher levels of inducible DNA damage whether we included Illinois turtles or not (Table 4, Fig. 4). Additionally, populations differed in their repair efficiency (Table 4). Analyses of the repeated levels of DNA damage across treatments (i.e., baseline, damage, and repair) revealed considerable variation among poplulations and sexes (Table S2; Fig. S1).

**Figure 4.**
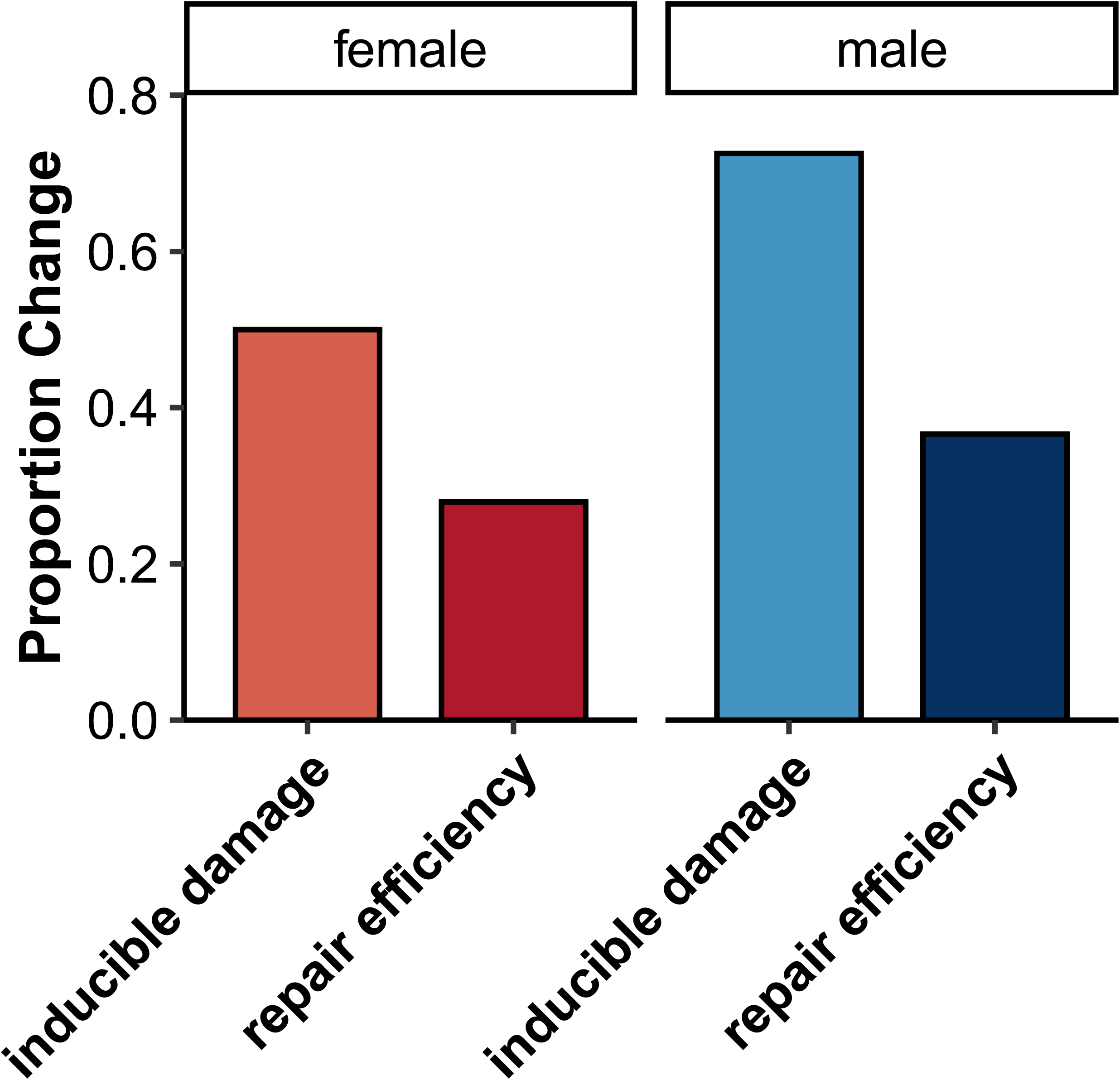
Visualization of the effect of sex differences in UV-inducible DNA damage and repair efficiency in erythrocytes across three populations of painted turtles. Values are back transformed LSmeans from the model in Table 4. Only inducible damage differed between males and females. Repair efficiency was not estimable for Illinois painted turtles.

**Table 4.** Analysis of variance of logit Inducible Damage and Repair Efficiency. Effect is the proportion of variation explained for explanatory variables with Pr < 0.15. NS is Pr > 0.20. zPl is z-transformed plastron length for each sex-by-population grouping to remove its association with both sex and population. D is Damage, B is Baseline, R is repair. Each of D, B, and R are proportion DNA in the tail of comets versus that in the head of the comet. See text for details.

|  | **Inducible Damage (logit)**<br>(D-B)/D |  | **Repair Efficiency (logit)**<br>(D-R)/B |  |
| --- | --- | --- | --- | --- |
| **Source of Variation** | **F<sub>df1, df2</sub>**<br>**(Pr > F)** | **Effect** | **F<sub>df1, df2</sub>**<br>**(Pr > F)** | **Effect** |
| Pop | 147.11<sub>2,36</sub><br>($<0.0001$) | 0.82 | 7.01<sub>1,21</sub><br>(0.015) | 0.18 |
| Sex | 5.67<sub>1,36</sub><br>(0.008) | 0.02 | 0.28<sub>1,23</sub><br>(NS) | -- |
| zPL | 0.40<sub>1,36</sub><br>(NS) | -- | 1.38<sub>1,21</sub><br>(NS) | -- |
| Sex x Pop | 0.73<sub>2,36</sub><br>(0.11) | -- | 0<sub>1,21</sub><br>(NS) | -- |
| zPL x Pop | 1.11<sub>2,36</sub><br>(NS) | -- | 0.08<sub>1,21</sub><br>(NS) | -- |
| zPL x Sex | 0.15<sub>1,36</sub><br>(NS) | -- | 0.61<sub>1,21</sub><br>(NS) | -- |
| ZPLxSex xPop | 3.85<sub>2,36</sub><br>(0.03) | 0.02 | 0.06<sub>1,21</sub><br>(NS) | -- |

#### Leukocyte differentials

We found significant variation among population-by-sex combinations for lymphocytes, basophils, and monocytes (Table 5). The effect sizes for this interaction, but no effect of zPL or the interactions with zPL (as a proxy for age) The magnitude of variation among populations was quite small for each type of cell (Table S3) Though not a focus of our study, the ratio of heterophils to lymphocytes, often used a proxy for immune stress (Holden et al. 2022), differed among populations (*F* _2, 117_ = 70, *Pr* < 0.0001) and between males and females (*F* _1, 117_ = 2.81, *Pr* < 0.10).

**Table 5.** Analysis of variance for four classes of leukocytes (ln-transormed values). Sex is male and female; zPl is z-transformed plastron length. Data were log10 transformed to improve normality. Non-informative interactions were removed.

|  | **Lymphocytes (ln)** |  | **Heterophils (ln)** |  | **Basophils (ln)** |  | **Monocytes (ln)** |  |
| --- | --- | --- | --- | --- | --- | --- | --- | --- |
| **Source of Variation** | **F<sub>df1, df2</sub>**<br>**(Pr > F)** | **Effect** | **F<sub>df1, df2</sub>**<br>**(Pr > F)** | **Effect** | **F<sub>df1, df2</sub>**<br>**(Pr > F)** | **Effect** | **F<sub>df1, df2</sub>**<br>**(Pr > F)** | **Effect** |
| Pop | 134.32<sub>2,120</sub><br>(<0.0001) | 0.66 | 35.23<sub>2,120</sub><br>(<0.0001) | 0.37 | 44.07<sub>2,119</sub><br>(<0.0001) | 0.39 | 18.84<sub>2,120</sub><br>(<0.0001) | 0.22 |
| Sex | 4.73<sub>1,120</sub><br>(0.03) | 0.02 | 1.46<sub>1,120</sub><br>(NS) | -- | 7.55<sub>1,119</sub><br>(0.007) | 0.06 | 0.19<sub>1,120</sub><br>(NS) | -- |
| zPl | 0.04<sub>1,116</sub><br>(NS) | -- | 0.22<sub>1,120</sub><br>(NS) | -- | 0.67<sub>1,119</sub><br>(NS) | -- | 0.18<sub>1,120</sub><br>(NS) | -- |
| Pop x Sex | 7.72<sub>2,116</sub><br>(0.0007) | 0.04 | 0.17<sub>2,120</sub><br>(NS) | -- | 4.43<sub>2,119</sub><br>(0.01) | 0.04 | 6.38<sub>2,120</sub><br>(0.002) | 0.07 |

## Discussion

In the past 25 years, López-Otín and colleagues (2013, 2023) proposed several biological hallmarks of aging that individually, or in concert, cause deteriorating function and accelerating mortality across the adult lifespan. Relatively few studies beyond those on model genetic species have sought to simultaneously measure cellular aging metrics across disparate hallmarks. However, we suggest that approaches using multiple metrics in the same individuals collected at the same time can yield insights into the evolution of aging mechanisms across diverse species. Such phenotypes in populations of wild species can provide imperative information on whether or to what extent these mechanisms are fixed or plastic. Here, we test two axes of intraspecific variation of interest – variation between the sexes and variation across populations in different types of cellular aging measures.

Two related hypotheses propose that sex-specific genomes contribute to aging. First, the “toxic Y” hypothesis posits deleterious mutation accumulation on the degenerate chromosome, impacting survival of the sex with that chromosome (Nguyen and Bachtrog, 2021). The other is the “unguarded X” hypothesis, which hypothesizes health and survival costs to the heterogametic sex due to the expression of deleterious mutations on the non-degenerate chromosome (Posynick and Brown, 2019). These genome architecture differences between males and females occur in many species. The presence of sex chromosomes in general have made the discovery of the causes of sex-biased aging and longevity difficult to separate from such differences in genome architecture. Species with environmental sex determination, such as many turtles, offer an opportunity to study sex differences in life histories between sexes with equal gene content and non-sex-linked genes. Painted turtles have sexual size, maturation, and survival dimorphism: females mature later, grow to a larger adult asymptotic size, and live longer than males (Warner et al. 2016; Hoekstra et al. 2018; Bronikowski et al. 2022). Thus, we predicted that males would show faster deterioration of cellular aging markers with advancing age than females.

In this study we addressed three general questions: (i) do measures of mitochondrial health – cellular metabolism and cellular reactive oxygen species – decline with age and/or size in a sex-specific manner; (ii) are measures of DNA damage affected by age and/or size in a sex-specific manner; and (iii) are age- and sex-specific patterns of mitochondrial health and DNA damage similar across a geographic context. We measured cellular basal metabolic rate, cellular maximal metabolic rate, cellular spare respiratory potential in PBMCs and reactive oxygen species concentrations, and measures of DNA damage in erythrocytes. We found that both sex and age can significantly impact these cellular hallmarks, but the pattern of this interaction varied across populations. Across all measures, sex had a much stronger effect than age/size (or its interaction with sex). Thus, these questions and their answers reveal complex and context-dependent mitochondrial health and DNA damage viz. aging. They suggest that although mitochondrial dysfunction and altered DNA repair are well-established hallmarks of aging (Brand and Nicholls, 2011; Jové et al. 2023; López-Otín et al. 2023), there is no general universal rule that declining bioenergetics and efficiency of mitochondria and increasing age-associated susceptibility to DNA damage are signatures of advancing age. We discuss these results in the context of observations from model species, other non-traditional species, and other species with large disparities in sex-related longevity.

### Mitochondrial “Dysfunction” as a hallmark of aging

#### Mitochondrial function

Age-related declines in cellular basal metabolic rate (Basal OCR) are well-documented in model species, such as mice *(Mus musculus*; Yuan et al., 2011; Bresilla et al. 2022), fruit flies (*Drosophila melanogaster*; Hoffman et al., 2014; Brown et al., 2020), and nematodes (*Caenorhabditis elegans*; Olsen et al, 2006; Hotzi et al., 2018), in studies that are typically performed within a biomedical context. Across a broader comparative context, studies that have tested age-related declines in mitochondrial function related to cellular metabolism generally find agreement with this decline, albeit via different mechanisms (humans (*Homo sapiens*) - increasing mtDNA mutations with age in both sexes (Short et al., 2005); garter snakes (*Thamnophis elegans*) - oxygen consumption rate declines with age in both sexes in slow aging ecotype (Gangloff et al., 2020); turquoise killifish (*Nothobranchius furzeri*) - overall decline in mitochondrial function in both sexes with age via mechanisms such as downregulation of mtDNA-associated genes (Hartmann et al., 2011)). Exceptions exist, however. For example, naked mole rats (*Heterocephalus glaber)* maintain exceptional mitochondrial health throughout their lifetime via the capacity of their mitochondria to consume damaging chemicals (H_2_O_2_) at significantly higher rates than similarly sized animals such as mice (*Mus musculus*; Munro et al., 2019). In our study, mitochondrial health, measured as spare respiratory potential, marginally increased with age in our Illinois population, especially in femals (Table 1). Spare respiratory potential is of interest, as it is analogous to respiratory reserve, which tends to decline with age (Desler et al., 2012). Indeed, a significant decline in Spare is a signature of aging in animals such as humans and mice, as rapid energetic requirements (due to environmental conditions, for example) cannot be met, leading to age-related diseases in a multitude of tissues (heart, lung, brain, etc.; Desler et al., 2012; Marchetti et al., 2020).

To test whether sex-by-age relationships in mitochondrial function are fixed or plastic across populations, we used body size as a proxy for age, as it scales with age in painted turtles (see Hoekstra et al., 2018). We found a significant three-way interaction between population (Idaho, Wisconsin, and Illinois), sex, and size (z-transformed plastron length) on all three measurements of mitochondrial oxygen consumption (Basal OCR, Max OCR and Spare; Table 3). Still, across these populations, males always had higher Basal OCR, and Max OCR, and Spare (Figure 3), suggesting population heterogeneity in the magnitude of how sex affects cellular respiration. Although we did not find consistent results supporting our prediction that Basal OCR would decrease faster in the faster-aging sex (males), it is possible that population differences in life histories exist, which could influence this trait in a manner different from what was predicted. These significant population interactions could also result from environmental differences during development across the three localities such as in Japanese quail (*Coturnix japonica*; Stier et al. 2022), though Bodensteiner et al. (2023) found remarkably homogeneous nest temperatures across painted turtle populations.

Life history-specific variation in resting metabolic rate could also exist across populations, as has been found in different species. For example, low food availability and potentially hypoxic conditions result in lower basal metabolic rates in carp (*Carpio carpio*; Killen et al., 2016). Fish in environments that require increased locomotion, and therefore have greater muscle density, have physiologies supporting higher Basal OCR (Killen et al., 2016). Life history trade-offs between Basal OCR and growth also occur in garter snakes (*Thamnophis elegans*) that exhibit either a slow or fast life history. Faster-growing ecotypes exposed to colder temperatures have significantly higher resting metabolic rates compared to the slow-growing ecotype (Gangloff et al., 2015). The turtle populations in the present study are not characterized as different ecotypes, but variation in ecological pressures may exist across the localities and could promote the observed phenotypic variation in Basal OCR rates. Changes in temperature, resource availability, etc., can impact basal metabolic rate and in different ways, depending on adaptive capabilities of the population (such as the case in rufous collared sparrows (*Zonotrichia capensis*; Cavieres and Sabat, 2008; Norin and Metcalfe, 2019).

The significant differences in mitochondrial health with sex and size in and across populations could indicate that the way mitochondrial health declines may not be universal. For example, mitochondrial density can differ between cell types (Faitg et al., 2021). Because we measured oxygen consumption rate in a heterogenous population of white blood cells rather than isolating mitochondria, cell type differences in mitochondrial density may have contributed, in part, to sex and age-differences. These mitochondrial differences can also vary across populations, such as in Atlantic killifish (*Fundulus heteroclitus*; Bryant et al., 2018; Chung et al., 2018; Hood et al., 2018). Mitochondrial physiology of hepatic origin differs across populations, but this pattern was not detected in other tissue types such as brain or heart (Chung et al., 2018; Hood et al., 2018). Other features of mitochondria, such as increases in mitochondrial variants within an individual (heteroplasmy), can lead to variation in mitochondrial function and contribute to aging phenotypes, particularly in males (e.g., fruit flies, Camus et al. 2012), although such sex-specific effects of heteroplasmy on phenotypes is not universal (e.g., greater mouse-eared bats (*Myotis myotis*), Jebb et al. 2018).

#### Mitochondrial reactive oxygen species

Cells have multiple ways of clearing damaging ROS (such as through enzymes superoxide dismutase, catalase, and other antioxidants such as glutathione and vitamin B). Aging-related increases in damaging free radicals are due to two complementary phenomena: decreasing efficiency of the electron transport chain with age and decreasing cellular abilities to neutralize ROS (Brand and Nicholls, 2011; Gómez et al., 2023). ROS in the context of damage and cell signaling have been studied across multiple taxa, acting both as damaging by-products of mitochondrial function and as signaling molecules modulating stress responses and longevity pathways (Shields et al. 2021). ROS are known to damage lipid membranes (Stark et al., 2005), DNA (Valko et al., 2004), and proteins (Stadtman, 2006).

In the context of aging, studies on the long-lived ocean bivalve clam *Artica islandica* have detected significantly lower levels of H_2_O_2_ production in comparison to the shorter-lived *Mercenaria mercenaria,* a closely-related mollusk (Ungvari et al. 2011). Furthermore, *A. islandica* removes this ROS faster than other related bivalves (Munro et al., 2023). Similarly, a comparative study among long-lived and short-lived colubrid snakes showed that the former exhibit lower hydrogen peroxide production than shorter-lived species (Robert et al., 2007). Conversely, long-lived western terrestrial garter snakes (*Thamnophis elegans*) have higher levels of ROS production relative to a co-occurring short-lived ecotype, which could indicate the signaling potential of these molecules to activate different cellular protection mechanisms under normal conditions (Schwartz and Bronikowski, 2012).

In turtles, it is hypothesized that their longevity is related to their ability to resist oxidative damage, among other factors. Freshwater turtles of the *Trachemys* and *Chrysemys* genera are facultative anaerobes, surviving long periods of hibernation under water. To prevent oxidative damage related to reoxygenation, these turtles have evolved several different antioxidant mechanisms. For example, *Trachremys scripta* have high constitutive levels of antioxidant enzymes, such as catalase, superoxide dismutase (SOD) and alkyl hyperoxide reductase (Willmore et al., 1997, reviewed in Krivoruchko and Storey, 2010).

In our study, we found a significant interaction between sex and age on mitochondrial superoxide (O_2_-) and intracellular hydrogen peroxide (H_2_O_2_), with young males having higher ROS levels overall. Female ROS levels were not affected by age, but male measures decreased with age (Fig 2a). Hoekstra et al. (2019) posit that expression of antioxidants from nucleated erythrocytes in reptiles may provide additional protection against ROS, yet the significantly lower levels of mitochondrial (O_2_-) and intracellular (H_2_O_2_) in aged males could explain, in part, the sex-specific difference in lifespan among painted turtles. Future studies could measure expression levels of superoxide dismutase, glutathione reductase, and other antioxidant enzymes to test whether there is concomitant increased antioxidant expression or activity (Holtze et al., 2021), facilitating the decrease in ROS levels, or if older males simply produce less of these oxidizing agents. High expression levels of Hsp60, a mitochondria chaperone that responds to oxidative stress, in turtles could also be involved in dampening the damaging effect of ROS in turtles (Chang et al., 2000, reviewed in Krivoruchko and Storey, 2010).

### “Genomic Instability” as a hallmark of aging

For our proxy measure for genomic instability, we measured inducible DNA damage and related repair efficiency (Aguilera and García-Muse, 2013). Specifically, we measured the ability of UVB (312nm) radiation to damage DNA, measured as the change between treated and untreated erythrocytes (using 5 minutes of UV exposure). UVB radiation causes adduct formation and base damage, which can lead to mutations and single-strand DNA breaks (Lehmann et al., 1998; Krokan and Bjoras, 2013). Along with irradiation, other genotoxic substances naturally existing in cells, such as ROS, can also cause single or double-strand breaks (Benhusein et al., 2010). Though we did not test different types of DNA damage, we found overall that sex significantly impacted inducible damage (i.e., the difference in DNA in the comet tail between the UV damaged cells and baseline damage; Table 4; Fig 4). We also found a significant effect of population on inducible damage and repair efficiency (Table 4). Long-lived animals, such as garter snakes (*Thamnophis elegans*), exhibit increased DNA repair mechanisms compared to similar shorter-lived species (Bronikowski, 2008) due in part to gene upregulation rates and variation in the number of genes implicated in the DNA repair process (Tian et al., 2017). Our study on painted turtles demonstrated intraspecific variation in DNA repair efficiency, indicating that there is either phenotypic plasticity or local adaptation in this trait. This is an important finding because enhanced DNA repair mechanisms are a signature of long lifespan, but our study shows that ecological differences can influence this trait. Indeed, diet has a significant impact on DNA damage in humans, where those with a higher antioxidant laden diet showed lower rates of strand breaks (Slyskova et al., 2014). Although this study did not show a direct relationship between diet and DNA repair efficiency, it is possible that population variation in environmental factors such as predation, diet, canopy coverage, etc., could influence this biomarker of aging in turtles, explaining the population differences detected (Ackerman and Horton, 2018).

In accordance with aging theory, we hypothesized that age would affect DNA damage and made the specific predictions that younger animals would have the lowest levels of baseline DNA damage and inducible DNA damage, and the highest DNA repair efficiency. We also predicted this would be particularly notable in females, which can live twice as long as males. Our results did not support this hypothesis. We found that young male turtles had lower levels of baseline damage compared to similarly aged females (Table 2), and this result was seen across populations when we used size as a proxy for age. At the same time, these results did support the prediction that inducible damage would be greater in the faster aging sex (males). The process of maturation, rather than age itself, may play a larger role in this mechanism, as an earlier study in painted turtles found that juvenile turtles had greater inducible UV damage than adults (Schwanz et al., 2011).

In conclusion, we found capricious variation in measures with age/size of mitochondrial function using modes of cellular oxygen consumption as our variables of interest. In contrast, the differences between males and females were strong and in agreement across populations – a finding that is particularly interesting when considering that genome content is the same between sexes in this species. This finding is important for understanding the basis of sex-biased longevity and aging in species lacking genotypic sex determination, and the relative contributions of age and sex to this finding. A promising future direction will be to focus on other aspects of mitochondrial health, such as reactive oxygen species production rates (e.g., Robert and Bronikowski, 2010), mitochondrial genome copy number, and expanded mitochondrial DNA damage quantification in young and old animals because mitochondrial dysfunction as a hallmark of aging encompasses numerous aspects of mitochondria – not all of which may be declining with age. Additionally, future research could focus on comparing mitochondrial functions across tissue types, which can also profoundly differ. For example, in rats, age-related activity in the electron transport chain differs across organs such as the brain and heart (Cocco et al., 2005). Bioenergetic measurements are only one component of understanding mitochondrial dysfunction. This biomarker is highly complex, so future avenues of research could also include testing decreased mitophagic responses (López-Otin et al., 2013), which is an autophagic response to remove damaged mitochondria and maintain mitochondrial homeostasis (Ding and Yin, 2012; Bakula and Scheibye-Knudsen, 2020; Chen et al., 2020). This degradation and recycling process is also subject to declining activity with age and, in some cases, in a sex-specific manner (Drummond et al., 2014; Levine and Kroemer, 2019; Bakula and Scheibye-Knudsen, 2020), therefore presenting itself as a fruitful direction for further investigation into mitochondrial dysfunction. Elsewhere, we have argued for greater inclusion of reptiles in comparative studies of aging physiology (Hoekstra et al. 2019; Reinke et al., 2022) to help place the voluminous mammalian and bird studies in an integrative amniote perspective on aging biology. Moreover, Nolin and Metcalfe (2019) highlight the need for more studies in natural populations in addition to lab conditions, where animals are necessarily at sub-optimal feeding levels, body temperatures, and mate seeking. Here we used a unique species with temperature-dependent sex determination in multiple wild populations to bridge this gap. These environmental factors can greatly impact components of metabolism and cellular repair mechanisms (e.g., Holden et al., 2021). Future studies should aim to further incorporate species with varied life history strategies because they can uniquely address questions on the ubiquitous nature of aging and its mechanisms. Understanding the ubiquity of cellular aging hallmarks is essential for an understanding of the complete life-history of species, sex differences in the aging process, and how these complete ontogenies respond to environmental change.

## Data Availability

Upon acceptance, raw data and scripts will be made available on Dryad.

## Acknowledgements

We thank A. Shortridge and V. Lacroix for their support with sample collection and leukocyte protocol establishment, respectively. And we thank L. P. Decena Segarro for help establishing the reactive oxygen species protocol. This research was supported by the following grants: NSF DBI-2213824 (AMB); NIH-NIA R01AG049416 (AMB, FJJ); NSF IOS-1257857 (FJJ); NSF BRC-BIO 2233233 (BAR). OA was an NSF REU in the Biology Integration Institute REU program supported by DBI-2213824 (AMB).

---

**Table S1.** Morphometrics for painted turtles from three populations (plastron length (PL), body mass (grams), ranges and sample sizes). In all cases, females are longer and weigh more than males (*Pr* < 0.05)

| Population | Female PL (mm)<br>(Mean ± SD,<br>range) | Male PL (mm)<br>(Mean ± SD,<br>range) | Female mass (g)<br>(Mean ± SD,<br>range) | Male mass (g)<br>(Mean ± SD,<br>range) |
| --- | --- | --- | --- | --- |
| Idaho<br>(n=30f, 17m) | 157±21<br>(127.5 to 195) | 128±17,<br>103.1 to 165 | 562±227,<br>265 to 1100 | 326±134,<br>175 to 610 |
| Wisconsin<br>(n=16f, 16m) | 147.1±19,<br>104.5 to 178 | 116.8±23,<br>83.2 to 150.2 | 556.6±173,<br>220 to 790 | 304.4±140,<br>130 to 500 |
| Illinois<br>(n=24f, 44m) | 163.7±9,<br>151.5 to 185.5 | 129.9±13,<br>97.1 to 159.7 | 751.2±121,<br>580 to 980 | 339.8±89,<br>170 to 500 |

**Table S2.** Counts of leukocyte types across populations and sexes for four classes of leukocytes. PBMCs are primarily lyphocytes and monocytes. Values are back-transformed LSmeans ln-transformed counts (see Table 5).

| Population | Lymphocyte | Heterophil | Basophil | Monocyte |
| --- | --- | --- | --- | --- |
| Idaho<br>Female<br>Male | 37<br>36 | 36<br>35 | 13<br>13 | 11<br>15 |
| Wisconsin<br>Female<br>Male | 38<br>40 | 25<br>23 | 24<br>20 | 13<br>16 |
| Illinois<br>Female<br>Male | 51<br>60 | 24<br>22 | 12<br>9 | 9<br>7 |

**Table S3.**
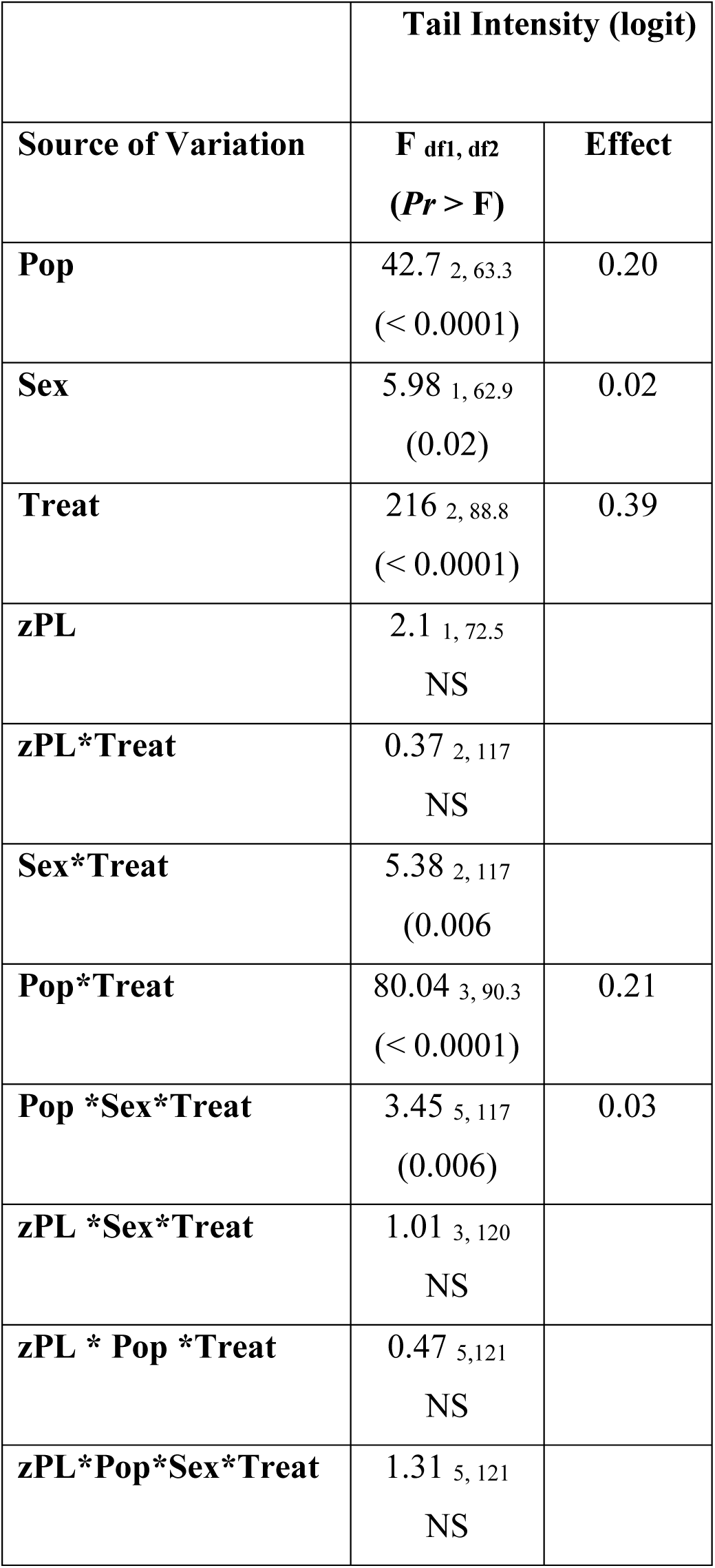
Repeated measures analysis of covariance for Percent DNA in comet tail under baseline, UV damage, and repair conditions. Treat is Treatment (B, D, R). Pop is Population. The (Sex x Treatment) interaction can be seen in Figure 4b. All data were logit transformed.

**Figure S1.**
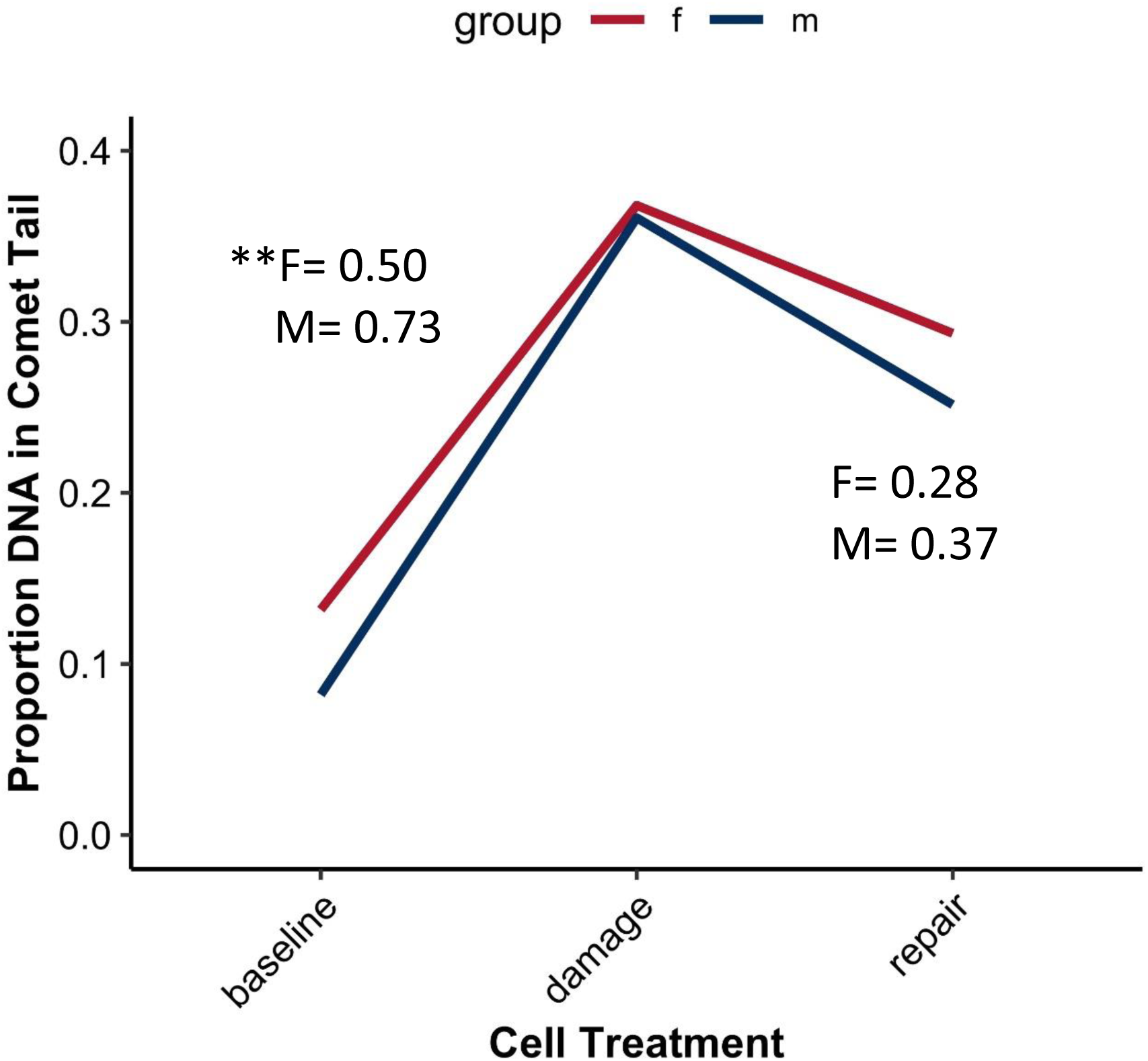
Visualization of the effect of sex differences in UV-inducible DNA damage in erythrocytes. Data are back-transformed least-square means of proportion DNA in the tail of comets versus the heads of comets from repeated measures analysis of variance (see Table S1) for baseline (B), UV-damaged (D), and repair (R). Values for inducible damage (%) and Repair efficiency (%) are overlaid. Only inducible damage differed between males and females. Repair efficiency was not estimable for Illinois painted turtles.

## Notes

### Competing Interest Statement

The authors have declared no competing interest.

